# Wavelength-encoded laser particles for massively-multiplexed cell tagging

**DOI:** 10.1101/465104

**Authors:** Nicola Martino, Sheldon J.J. Kwok, Andreas C. Liapis, Sarah Forward, Hoon Jang, Hwi-Min Kim, Sarah J. Wu, Jiamin Wu, Paul H. Dannenberg, Sun-Joo Jang, Yong-Hee Lee, Seok-Hyun Yun

## Abstract

Large-scale single-cell analyses have become increasingly important given the role of cellular heterogeneity in complex biological systems. However, no current techniques enable optical imaging of uniquely-tagged individual cells. Fluorescence-based approaches can only distinguish a handful of distinct cells or cell groups at a time because of spectral crosstalk between conventional fluorophores. Here we show a novel class of imaging probes emitting coherent laser light, called *laser particles*. Made of silica-coated semiconductor microcavities, these laser particles have single-mode emission over a broad range from 1170 to 1580 nm with sub-nm linewidths, enabling massive spectral multiplexing. We demonstrate the stability and biocompatibility of these probes *in vitro* and their utility for wavelength-multiplexed cell tagging and imaging. We demonstrate real-time tracking of thousands of individual cells in a 3D tumor model for several days showing different behavioral phenotypes. We expect laser particles will enable new approaches for single-cell analyses.

---

The emerging understanding of cellular heterogeneity in biology has necessitated new tools for single-cell analysis. Single-cell sequencing has helped define cell types with increasingly sophisticated details and revealed the critical role of cellular heterogeneity in cancer, stem cell differentiation, and epithelial homeostasis^1–4^. Despite these advances, a major challenge is the ability to tag and discriminate individual cells, and track them over time or over the course of different measurements. Current fluorescence-based approaches are limited in multiplexing ability, and are insufficient, for example, for visualizing the hundreds to thousands of distinct clonal lines present in tumors^1,5^. Biomolecular techniques using unique DNA barcodes can effectively label a virtually unlimited number of cells^3,6^; they are however incompatible with imaging and do not enable retention of spatial information. Furthermore, sequencing to read the DNA barcode is destructive, requiring cell dissociation and lysis, and thus limited to only endpoint measurements. These limitations highlight the need for unique tags with optical readout.

While fluorescence microscopy is widely used for cellular imaging^7^, typical fluorophores linewidths, between 30–100 nm, allow no more than a few spectra to fit in the entire visible spectrum without overlap. Consequently, this technique can normally resolve only a handful of labels, preventing concurrent study of many more cell types and subtypes. It is fundamentally challenging to engineer fluorophores or inorganic emitters to have much narrower emission spectra because of the quantum mechanical and thermodynamic broadening of their electronic energy levels. Raman emission from vibrational transitions is still broad (~20 nm)^8^. Multiple fluorescence emitters with dissimilar spectra can be combined^9–11^, but these approaches relying on intensity-based spectral analysis or nanoscale super resolution are of limited use for large-scale tracking in optically dense tissues due to wavelength-dependent absorption and scattering. As opposed to molecular engineering, photonic principles harnessing optical resonance and amplification could offer a solution. By placing fluorescent emitters inside an optical cavity with a sufficient Q-factor, laser emission with extremely narrow, sub-nanometer linewidth can be produced^12^. Previous efforts using dye-doped microspheres have shown the proof of concept of this photonic approach^13,14^, although the large resonator sizes of ~10 μm has prohibited practical applications.

Here, we show microlasers with massive multiplexing capability and optimized properties for cell tagging and tracking applications. We name these imaging probes *laser particles* (LPs). To miniaturize laser sizes, we used semiconductor materials with high refractive index^15^ and gain in a microdisk geometry. While microdisk lasers have been extensively investigated for on-chip applications, we developed methods to detach them from the substrate, suspend them in solution and coat them with a biocompatible and protective layer (Fig. 1a). These laser particles occupy only ~0.1% of a typical cell’s volume and generate single narrowband emission peaks (~0.4 nm), tunable across a wide spectral range of 400 nm. We show their stable performance in cells and present methods to identify tagged cells using laser particle stimulated emission (LASE) microscopy. Finally, we demonstrate massively multiplexed tracking of thousands individual cancer cells in a 3D tumor spheroid invasion assay. Our work establishes laser particles as a new class of luminescent probes that expands optical microscopy for large-scale, comprehensive single-cell analysis.

**Fig. 1.**
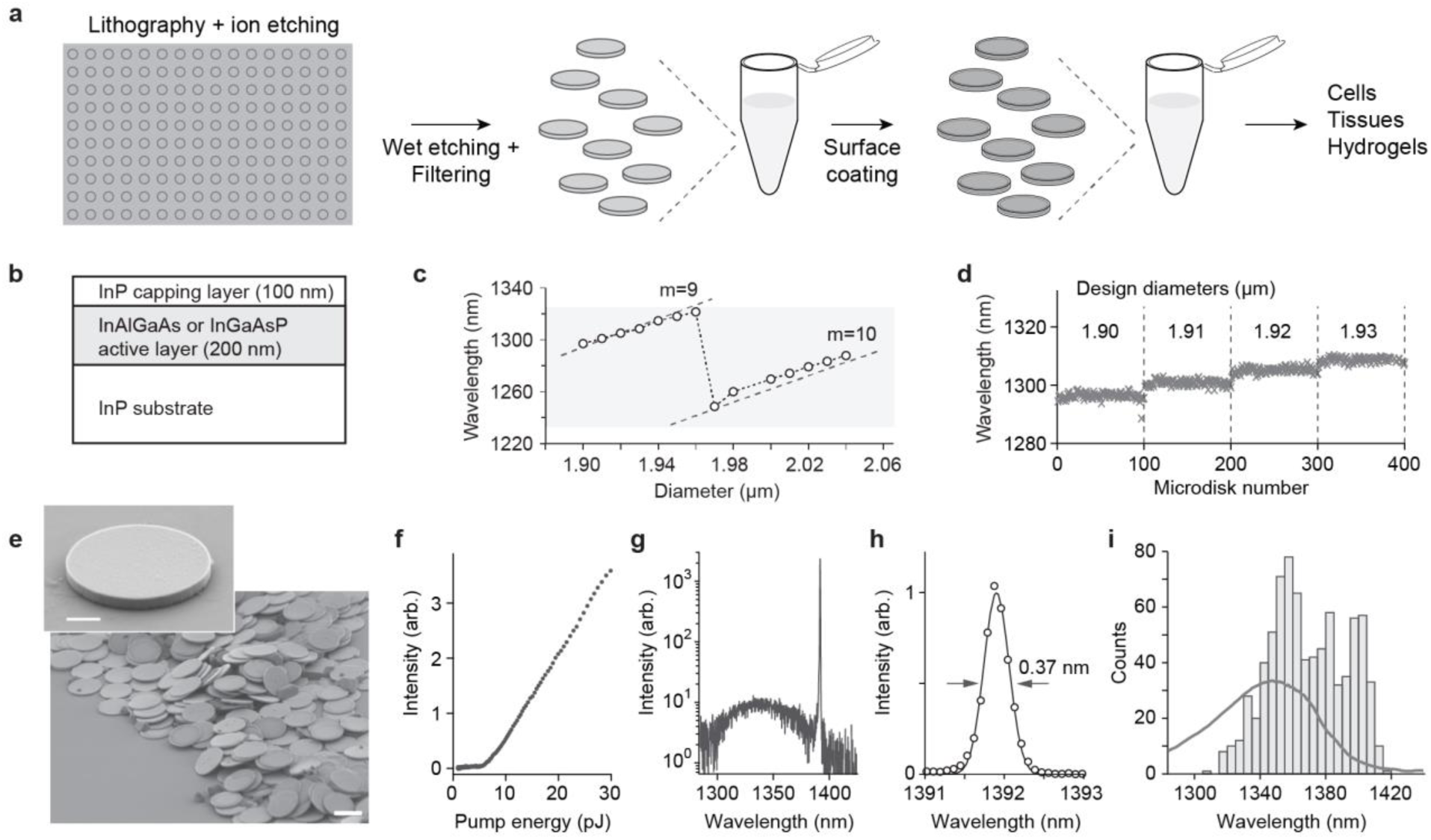
Optical properties of semiconductor microdisk lasers. **a**, Schematic representation of the production process of our laser particles. **b**, Structure of the epitaxial wafers used for the fabrication of microdisk lasers. c, Measured emission wavelength of microdisks (in air, on pillar) with increasing design diameters varying from 1.9 to 2.04 μm in steps of 10 nm. The shaded box corresponds to the gain region of the semiconductor (In_0.73_Ga_0.27_As_0.58_P_0.42_). The dashed lines are the calculated cavity-mode resonance wavelengths for a microdisk (in air) with a refractive index *n* = 3.445. **d**, Measured center wavelength of laser emission from microdisks on pillars. Four groups of samples fabricated with different design diameters in 10 nm steps are shown. The standard deviation (σ) is ~1 nm (*N* = 100). **e**, SEM image of microdisk resonators after detachment from the wafer (scale bar: 2 μm). The inset shows a close-up of a microdisk (scale bar: 500 nm). **f**, Output curve of laser emission versus pump energy for a typical cavity. **g**, Typical output emission spectrum of a microdisk resonator (*E_p_* = 20 pJ) above threshold. **h**, Gaussian fit of the laser emission peak. **i**, Histogram of the emission wavelengths of *N* = 794 different microdisk resonators suspended in Matrigel overlaid on the fluorescence spectrum of the active material (In_0.53_Al_0.13_Ga_0.34_As).

## Single-mode microlasers over a wide spectral range

Among several semiconductor materials suitable for laser particles, we chose InAlGaAs and InGaAsP quaternary alloys (Fig. 1b), which have small bandgap energy for operation in a near-infrared (NIR-II) region of 1.0–1.8 μm^16^. This range is attractive owing to low phototoxicity to cells, relatively good penetration in tissues, and no spectral overlap with conventional fluorescent probes^17,18^. We chose a microdisk design supporting planar whispery-gallery-mode (WGM) resonances^19^ because sufficient passive-cavity Q-factors of > 1,000 can be obtained with micron or submicron sizes^20^, and the resonance wavelength is tunable by changing its diameter^21^.

Calculations based on WGM theory (see Supplementary Note 1) predict that, for microdisks with diameters around 2 μm, the sensitivity of the resonance wavelength to small changes in diameter is around 1 nm/nm. Using electron(e)-beam lithography we produced batches of microdisks with identical design diameters separated by 10 nm steps. Measurements of their emission wavelengths closely followed theoretical predictions, with a discontinuity when the resonance jumps to a higher order as the diameter increases (Fig. 1c). Measurements of *N* = 100 disks of the same batch show a standard deviation σ = 1 nm in resonance wavelength (Fig. 1d). Although this spectral uniformity can be useful and may be further improved, in this work we chose to use UV lithography instead to be able to produce microdisks in large quantity (~3.2 million microdisks per cm^2^ of wafer) at much faster speed and lower cost than e-beam lithography. We allowed microdisk diameters to vary over a ~200 nm range so that the fabricated microdisks have randomly varying wavelengths over the entire gain bandwidth of the semiconductor. We tested different wafer designs including multi-quantum-well (MQW) structures and bulk semiconductor active layers and obtained comparable performance in terms of laser threshold and emission linewidth (Supplementary Fig. 1). The LPs described below were fabricated from bulk In_0.53_Al_x_Ga_0.47-x_As or In_x_Ga_1-x_As_y_P_1-y_ epitaxial layers.

Microdisks were released from the substrate via wet-etching in hydrochloric acid solution, and suspended in water after removing debris using size-selective filters. Scanning electron microscopy (SEM) showed reproducible microdisks with smooth edges (Fig. 1e). For optical characterization, the collected microdisks were embedded in 3D hydrogel matrices (Matrigel). Upon optical pumping with a pulsed Ytterbium-doped fiber laser, the output emission from each microdisk was analyzed using a NIR spectrometer. Lasing threshold was observed at a pump energy (*E_p_*) of ~7 pJ for a beam diameter of ~2 μm (fig. 1f). Above threshold, the output spectrum featured a single peak, with a full-width-at-half-maximum (FWHM) linewidth of ~0.4 nm (Fig. 1g, h). The measured intensity varied considerably depending on the microdisk orientation in the hydrogel because WGMs radiate predominantly in the radial direction. The lasing wavelengths of microdisks obtained from a single epitaxial wafer (In_0.53_Al_0.13_Ga_0.34_As on InP) spanned a broad region across 100 nm, supported by the gain bandwidth of the active medium (Fig. 1i).

To extend wavelength coverage, we used five semiconductor wafers with their fluorescence peaks separated by ~80 nm (Fig. 2a). We calculated WGM modes^22^ as a function of diameter (Supplementary Fig. 2) to confirm that, at a given diameter, only 1 or 2 cavity modes fall in the gain bandwidth of each wafer (Fig. 2b). Experimentally, microdisks of this diameter range generated single-mode laser emission under normal operation conditions. We were thus able to produce batches of microdisks with single-mode emission at different wavelengths across an ultrawide spectral range from 1170 to 1580 nm (Fig. 2c).

**Fig. 2.**
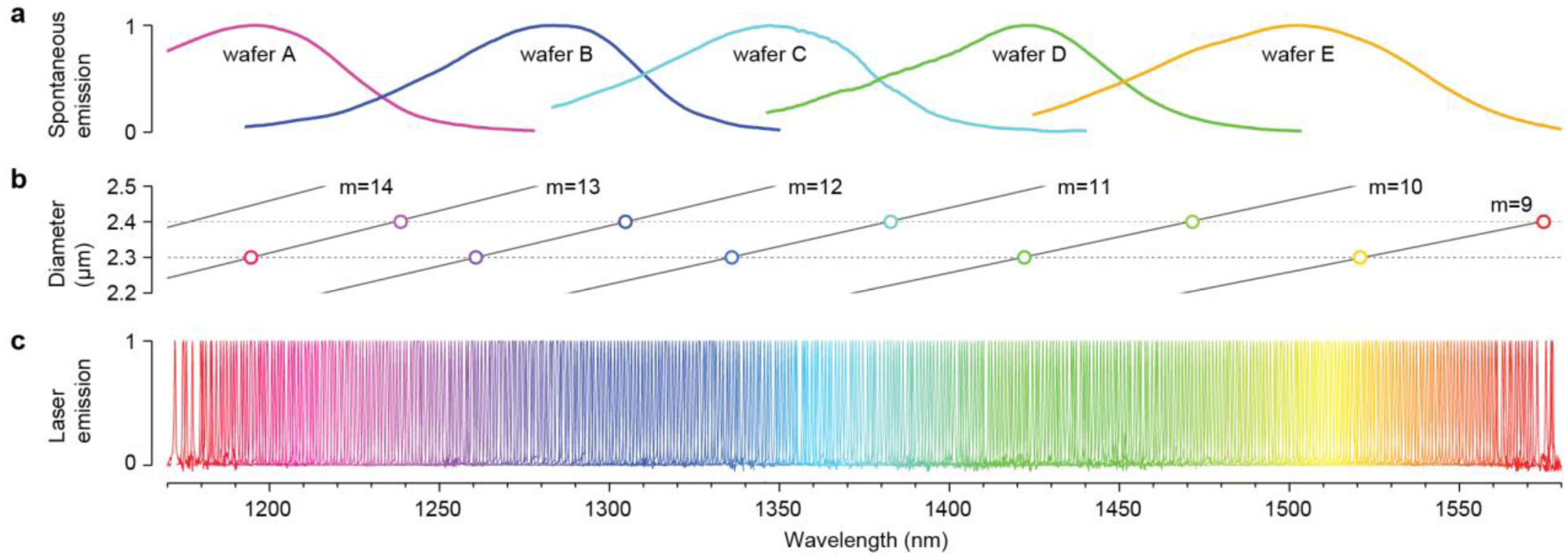
Highly multiplexed microdisk lasers. **a**, Photoluminescence (fluorescence) spectra of five different semiconductor materials used in this work; wafer A: In_0.80_Ga_0.20_As_0.44_P_0.56_, wafer B: In_0.73_Ga_0.27_As_0.58_P_0.42_, wafer C: In_0.53_Al_0.13_Ga_0.34_As, wafer D: In_0.53_Al_0.09_Ga_0.38_As, and wafer E: In_0.53_Ga_0.47_As_0.92_P_0.08_. **b**, Calculated resonance wavelengths of WGM modes with mode-order *m*, for different microdisk diameters between 2.2 to 2.5 μm. Circles represent possible lasing modes obtainable from the five different wafers with microdisk sizes of 2.3 and 2.4 μm, respectively. c, Normalized laser emission spectra of 400 laser particles in a range from 1170 to 1580 nm with an interval of ~1 nm. All laser particles were pumped by a common laser source.

## Silica coating of semiconductor microdisks

Unprotected non-oxide semiconductor materials tend to slowly corrode in water and, under photoexcitation, can generate undesirable electrochemical effects^23,24^. To enable operation in aqueous biological environments, we developed a protocol to passivate the semiconductor surface of the microdisks by coating with silicon dioxide (SiO_2_). Each cycle of a modified Stöber process^25^ produced a silica layer of ~50 nm, and multiple cycles resulted in thicker coating (Fig. 3a, b). Cross-sectional SEM, energy dispersive X-ray spectroscopy (EDS) and transmission electron microscopy (TEM) confirmed uniform silica coating (Fig. 3c, d, Supplementary Fig. 3).

**Fig. 3.**
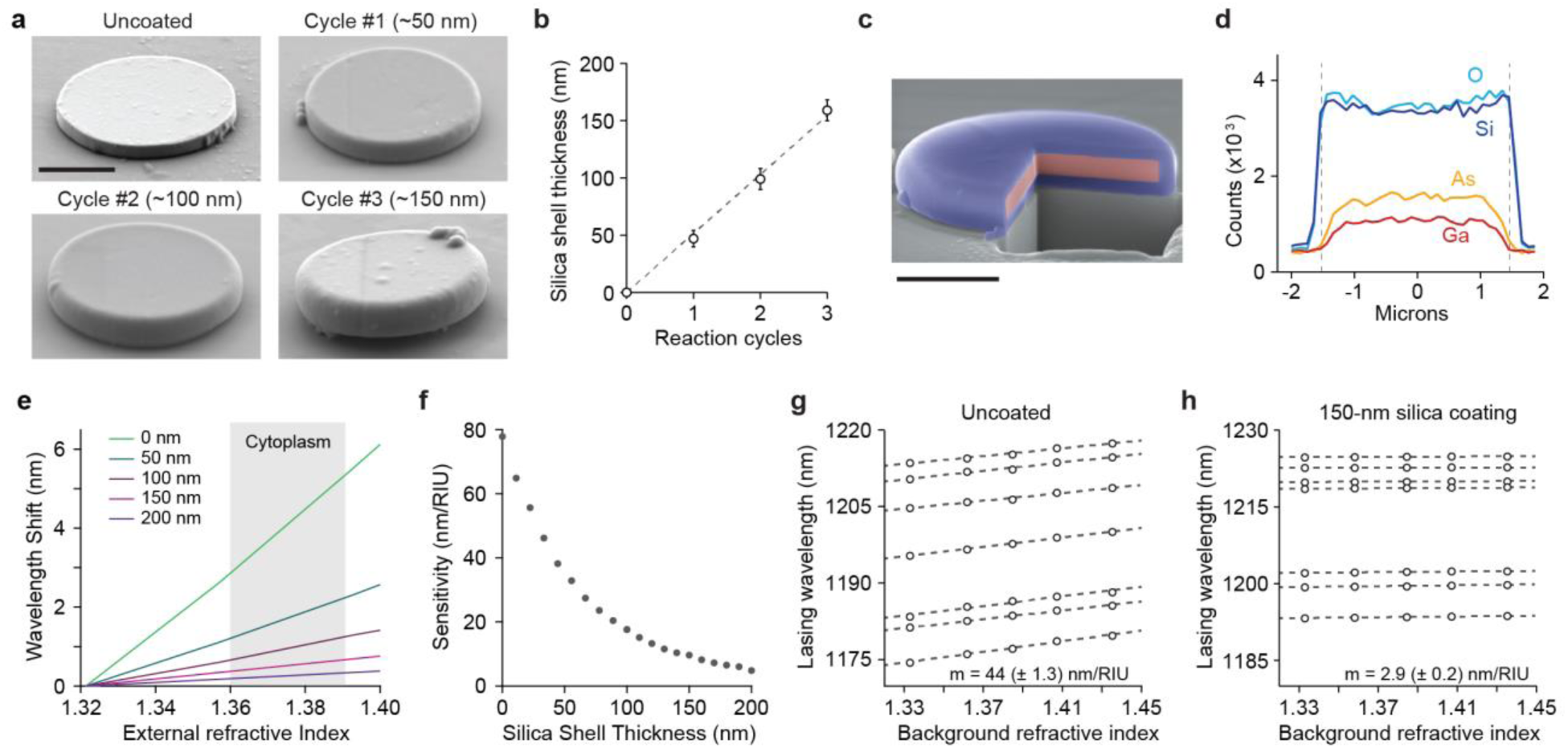
Silica coating of III-V semiconductor microdisk lasers. **a**, SEM images of microdisk resonators before and after 1,2 or 3 cycles of coating. Scale bar: 1 μm. **b**, Dependence of silica shell thickness on reaction cycles measured by TEM. Mean ± 95% CI are presented, with *N* ≥ 9 for each condition. **c**, False-color cross-sectional SEM image of a coated microdisk cut with focused ion beam (FIB). Scale bar: 1 μm. **d**, EDS analysis of different elemental species along the diameter of a coated microdisk. **d**, Measured laser emission wavelengths of a silica-coated (150 nm) microdisk and an uncoated microdisk placed on a glass substrate and immersed in sucrose solutions with different refractive indices. **e**, Wavelength shift of a microdisk resonator with respect to change in external refractive index, calculated from FDTD simulations for increasing thicknesses of silica coating. The grey shaded region corresponds to the typical range for the refractive index inside the cytoplasm. **f**, Sensitivity of the microdisk resonance wavelength to changes in external refractive index as a function of increasing silica coating thickness, calculated for small variations around the typical value in cytoplasm (*n_1_* = 1.37). **g**, **h**, Dependence of lasing wavelength on background refractive index for various uncoated (g) and 150 nm coated (h) microdisks. Measurements were taken from microdisks on a glass substrate immersed in sucrose solutions of different concentrations.

Besides material protection, another critical role of silica coating is decreasing the evanescent field of cavity modes in the surrounding medium and thereby reducing the sensitivity of lasing wavelengths to changes of external refractive index (*n*_ext_). Finite-difference time-domain (FDTD) calculations predict a wavelength dependence on refractive index (Δ*λ*/Δ*n*_ext_) of 80 nm/RIU for uncoated microdisks, which corresponds to a variation of up to 2.4 nm in cell cytoplasm (*n*_ext_ = 1.36–1.39)^26^, while this sensitivity decreases with increasing coating thickness (Fig. 3e, f). The wavelength sensitivity of 150-nm-coated microdisks deposited on a glass substrate was measured to be 2.9 ± 0.2 nm/RIU, much lower than the 44 ± 1 nm/RIU for uncoated microdisks (Fig. 3g, h).

## Stability of laser particles in biological environments

Our standard design coating thickness was 100 nm. Coated microdisks embedded in cell-culture hydrogels had slightly higher threshold pump energy of ~9 pJ compared to uncoated lasers in hydrogels (~7 pJ) due to the higher refractive index of silica (1.46) than of the hydrogel (1.34) (Fig. 4a). However, silica coating dramatically improved the optical stability of microdisks under continuous pulsed excitation (*E_p_* = 40 pJ). After 1 billion pulses, uncoated microdisks in hydrogels showed output power degradation by about 20%, and an exponential decrease in lasing wavelength of 1.5 nm, both of which were less stable than in air (Fig. 4b, c and Supplementary Fig. 4). The degradation of uncoated microdisks is attributed to surface oxidation^27^ and photochemical etching of the semiconductor^24^. According to our simulations (see Supplementary Note 1), a surface corrosion of the semiconductor by 2 nm causes a spectral blue-shift of ~1.8 nm. Silica-coated microdisks in hydrogels were much more stable, producing constant intensity and a minute wavelength shift of 0.1 nm after emitting 1 billion laser pulses. Hard silica coating greatly reduced the water-induced degradation of semiconductor materials. Over 30 hours in cell culture medium, coated samples showed stable lasing wavelengths within 0.4 nm, whereas uncoated semiconductor microdisks degraded with blue-shifts of 11 nm (Fig. 2d).

**Fig. 4.**
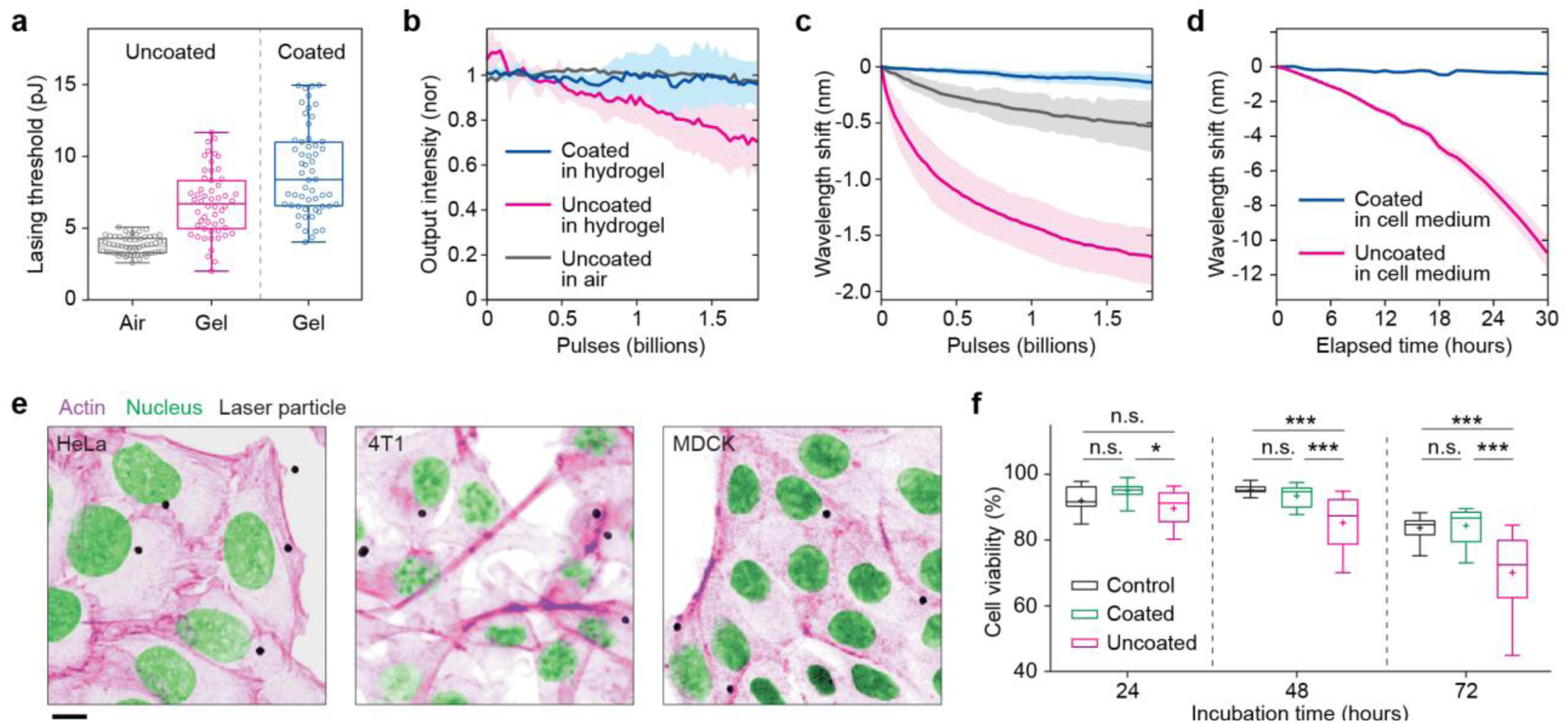
Stability and biocompatibility of laser particles. **a**, Distribution of lasing thresholds for uncoated microdisks in air (suspended on a small pillar before complete detachment from the substrate; (*N* = 60), uncoated microdisks in Matrigel (*N* = 54) and coated microdisks in Matrigel (*N* = 56). **b**, Lasing output intensity and **c**, resonance wavelength shift of uncoated microdisks in air, uncoated microdisks in Matrigel, and coated microdisks in Matrigel under continuous illumination up to 1.8 billion pump pulses (*E_p_* = 40 pJ/pulse, repetition rate 2 MHz); solid lines are mean (*N* = 4 per group) and shaded regions are 95% confidence intervals. **d**, Lasing-wavelength shifts of coated (*N* = 51) and uncoated microdisks *(N* = 6) on glass substrate in cell culture medium. Solid lines are mean and shaded regions are 95% confidence intervals. The samples were kept under darkness except during measurements taken every hour using a short exposure to pump pulses. **e**, Confocal fluorescence images of HeLa, 4T1 and MDCK-II cells with staining for actin (magenta), and nucleus (green), overlaid with brightfield transmission images of LPs (grayscale). Scale bar: 10 μm. **f**, Cell viability measured using a live/dead fluorescent assay at 24, 48 and 72 hours after incubation with LPs. MDCK-II cells were cultured in serum-free media with silica-coated or uncoated LPs at a particle-to-cell ratio of 1:1, and compared to no particles (control). A two-way ANOVA [F(2,22) = 16.47] for the effect of coating on cell viability was significant (p < 0.05) at 48 and 72 hours, ^∗^p < 0.01, ^∗∗∗^p < 0.001 using Tukey’s test.

The silica coating can be readily modified to provide additional functionalities to LPs, such as multimodal imaging, biomolecule sensing, or cell-type specific targeting^28–30^. For example, incorporation of fluorescent dye into the silica shell allows fluorescence-based detection of LPs, encapsulation of the silica surface with poly(ethylene glycol) can reduce non-specific cellular interactions^31^, and coating with biotin enables conjugation with biomolecules of interest (Supplementary Fig. 5).

## Biocompatibility of laser particles

The silica coating is also essential to improve biocompatibility of the laser particles. Several cell lines efficiently internalized LPs within 24 hours of incubation *in vitro* through the non-specific process of macropoinocytosis^29,32^ (Fig. 4e and Supplementary Video 1). A calcein AM/ethidium homodimer-1 fluorescent assay showed that while coated LPs had no measurable effect on cell viability compared to control, significant toxicity was induced by uncoated microdisks after incubation for 48 and 72 hours (Fig. 4f and Supplementary Fig. 6). This result is consistent with previous studies on silica-coated quantum dots, in which the coating layer prevents leakage of toxic ions from the semiconductor surface^33^.

To assess the suitability of our laser particles as intracellular probes, we performed time-lapse imaging to observe the migration and proliferation of MDCK-II cells containing LPs for several days (Supplementary Videos 2, 3). As cells divided, LPs were transmitted from a mother cell to daughter cells. Mitotic partitioning of multiple LPs tended to be asymmetric, fitting a skewed binomial probability distribution (Supplementary Fig. 7). In serum-free culture, in which cell division is minimized, LPs uptake followed a Poisson distribution (Supplementary Fig. 8), indicative of a stochastic process. For cells containing up to 6 particles, their presence did not significantly affect cell cycle times (Supplementary Fig. 8). A cell proliferation assay (CCK-8) also revealed no significant difference in proliferation rate between cells with and without LPs (Supplementary Fig. 8). A few rare cells with an excessive number (> 10) of particles were unable to complete mitosis and underwent apoptosis (Supplementary Video 3).

For long-term cell tracking, intracellular laser particle retention is important. Previous works have shown that for sub-micron particles the rate of exocytosis decreases with increasing particle size, irrespective of cell type^34^. Particles of diameters 1–3 μm were found to remain internalized in cells for at least 6 days^35,36^. We followed hundreds of LP-tagged MDCK-II cells *in vitro* using time-lapse imaging for over 60 hours (Supplementary Videos 1–3) and did not observe any clear evidence of exocytosis (Supplementary Video 4). While we cannot exclude the possibility of LPs being released after cell death, appropriate surface coatings may prevent their reuptake^37^.

## Imaging of intracellular laser particles

For high-speed imaging of laser particles, we coupled the pump laser and spectrometer to a laserscanning confocal microscope. The pump beam under-filled the objective lens (20X) such that the focal beam size was 2 μm, matching the size of LPs. The system can acquire the LASE images of LPs (with a pixel integration time of 100 μs), the brightfield transmission and confocal fluorescence images of cells and tissues. For example, LASE-fluorescence microscopy showed signatures of individual LPs with varying shapes depending on their orientation inside the cytoplasm of HEK-293 cells expressing membrane-localized green fluorescent protein (GFP) (Fig. 5a).

**Fig. 5.**
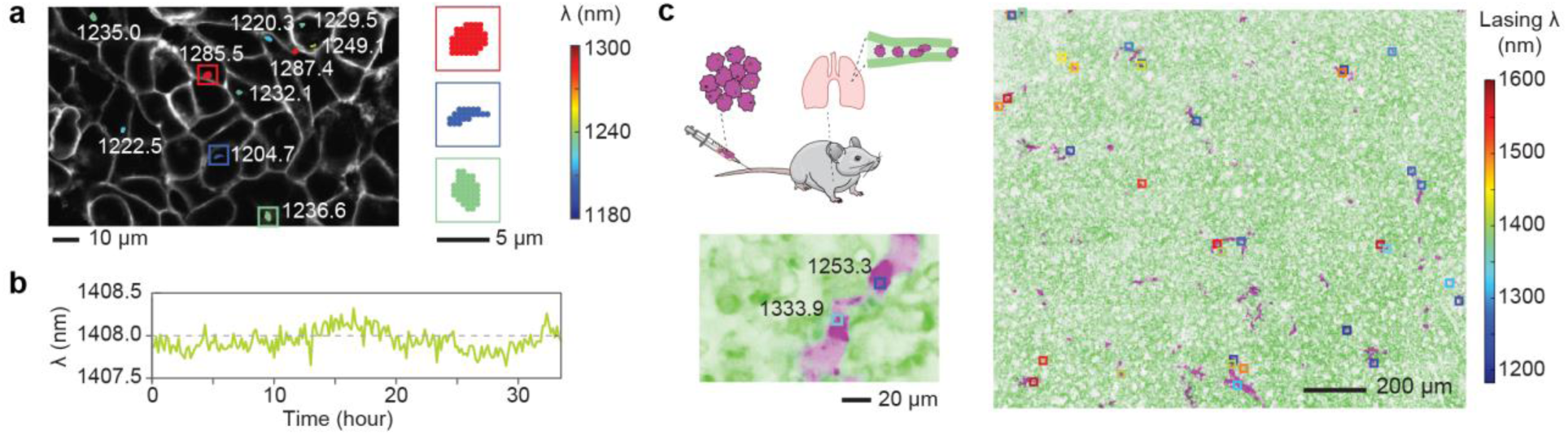
High-speed imaging and characterization of laser particles as cell tagging probes. **a**, Overlaid LASE-fluorescence imaging of LPs inside membrane-GFP-expressing HEK-293 cells. Inset, zoomed-in images of three LPs, in which the color of each dot (pixel) represents the peak wavelength of laser emission. **b**, Representative wavelength trace of an intracellular LP measured every 10 minutes for 33 hours. **c**, LP-tagged, fluorescence-dye-labeled 4T1 cells were injected intravenously into a live mouse. After 15 minutes, lung tissue was removed for *ex vivo* imaging. Maximum intensity z-projection images show administered 4T1 cells (purple), lung parenchyma (green), and the positions (squares) and wavelengths (color) of LPs in the lung.

Using a compact cell-culture incubator on the microscope, we acquired time-lapse LASE images of laser particles inside cells. Z-stack images were acquired at each time point, from which the position and wavelength of each LP were determined by using a clustering algorithm (Supplementary Fig. 9). The output wavelengths of LPs in the cytoplasm were stable (Fig. 5b), apart from random fluctuations (0.1 nm) and slower variations. These changes were attributed to the residual sensitivity to the surrounding refractive index that changes slowly by natural cellular processes and rapidly by random-walk movement of LPs inside the cytoplasm. The wavelength variation exhibited a normal distribution with standard deviation σ = 0.18 nm. For LPs with nominal difference in wavelength of Δ = 1 nm, this correspond to an error probability in identifying LPs of *P* ≈ 10^−4^. This error can be reduced by time-lapse spectral measurement averaging random fluctuations or adding redundancy such as positional information.

We then tested the possibility of tagging and imaging cells in tissues. 4T1 cells were stained with a fluorescent dye and loaded with LPs; they were then injected into the tail vein of a membrane-GFP expressing mouse, in a model mimicking hematogenous micrometastasis. After 15 minutes, the lungs (where most of the injected cells are trapped) were explanted and imaged with our LASE system. Co-localization of LP emission with the cellular dye demonstrates their reliability as intracellular tags also in scattering tissues (Fig. 5c).

## Longitudinal tracking in tumor spheroids

To demonstrate large-scale cell tracking, we used a 3D tumor spheroid model using LP-tagged polyclonal 4T1 breast cancer cells. A single spheroid contained about 70,000 LPs. Using the LASE microscope setup described above, it took 47 min to acquire a z-stack 3D scan over imaging volume of 1 x 1 x 0.28 mm^3^. A computer-automated 3D scan was conducted every hour for 128 hours during which the spheroid expanded in size (Fig. 6a and Supplementary Video 5). At each time point, 4,500 to 8,000 LPs were detected (Fig. 6b and Supplementary Fig. 10), with the number increasing over time as the tumor grew toward the glass bottom plate and more LP-tagged cells entered, while some left, the imaging volume. Because the multiplicity of our current batch of LPs is lower than the number of tagged cells, we used their positional information (geometrical proximity) together with the spectral information to minimize tracking error. This tracking algorithm enabled us to track 75–80% of all detected laser particles for longer than 24 hours, among which 731 were tracked for the more than 125 hours (Supplementary Fig. 10). We note that tracking fluorescently-labeled cells in 3D tissues would have to rely on purely geometrical information and would require much smaller time steps, which is technically prohibitive and prone to photobleaching. The 3D trajectories reveal a variety of migratory patterns (Fig. 6c). Long trajectories typically show several intermittent slow regions with an interval of about 1 day, which we hypothesize to result from cell division. We found groups of 2–3 LPs travelling in very close proximity, which we interpreted as being inside the same cells (Supplementary Fig. 10). Some of two initially co-traveling LPs separated into two distinct paths (Fig. 6d). This phenomenon is likely due to splitting of particles to different descendent cells during cell division.

**Fig. 6.**
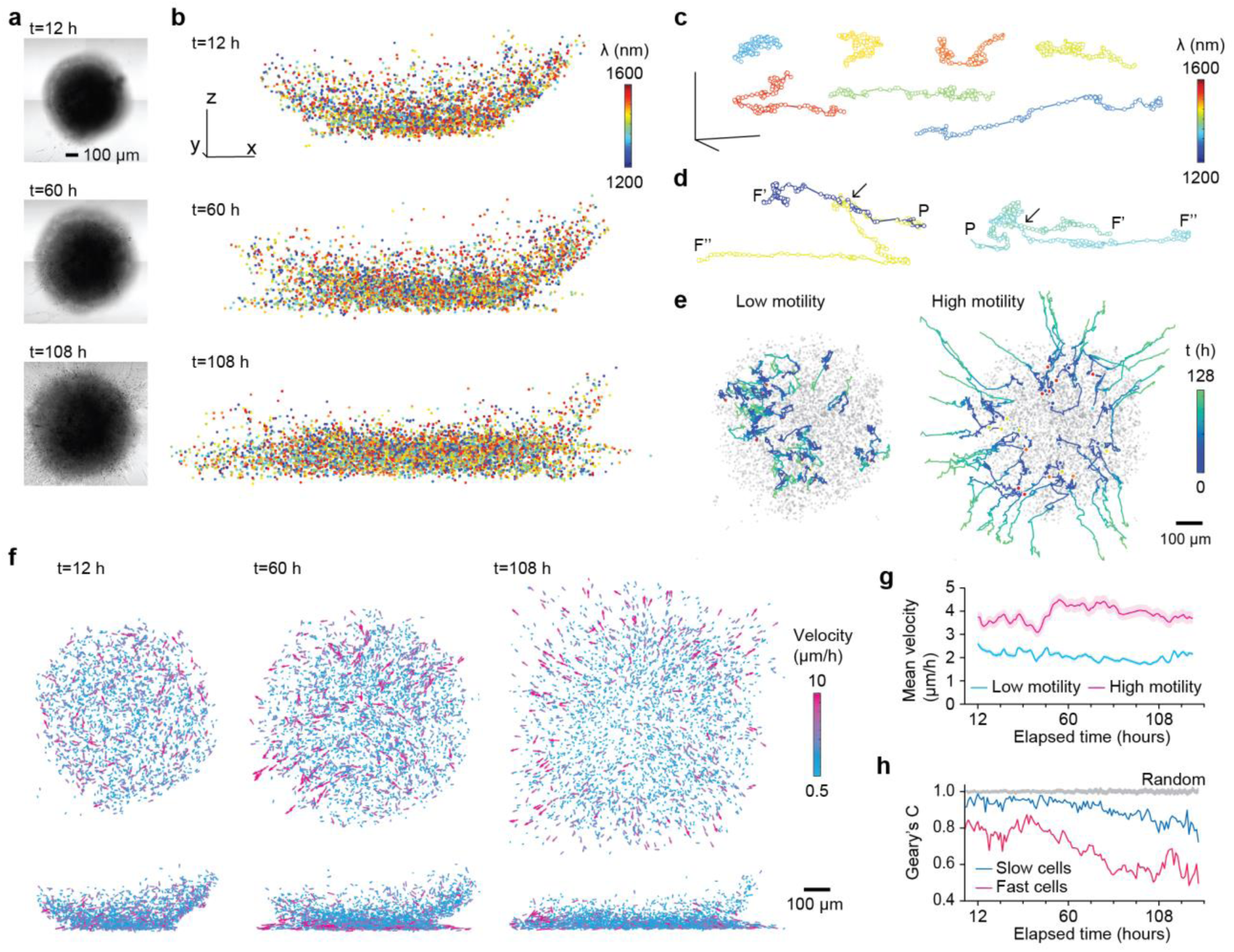
Cell tracking in a tumor spheroid. **a**, Optical transmission image of the tumor spheroid at 12, 60 and 108 h. **b**, Spatial distribution of LPs inside the tumor. Each dot represents a LP, with color coding for its wavelength. **c**, Trajectories of several exemplary LPs; circles indicate measured positions every hour. **d**, Trajectories of single parental cells (P) separating into two descent cells (F’ and F”) halfway during tracking (arrows). **e**, Representative paths of cells divided in two groups in terms of average motility: high (top 25%), and low (bottom 25%), projected in top view (XY). The initial position of each LP is marked with a circle, color-coded by wavelength; the blue-to-green color scheme in individual paths denotes elapsed time over 128 h. Grey dots denote the positions of all detected LPs at 12 h to visualize the overall spheroid shape. **f**, Maps of instantaneous velocities of tracked cells at different time points. **g**. Mean instantaneous velocities of cells in the high and low motility groups over time for cells tracked from 12 h to 120 h. *N* = 200 cells per group. Shaded region indicates 95% confidence interval. **h**. Geary’s coefficient for spatial correlation of cell velocities as a function of time for all tracked cells. Fast and slow cells represent the top 25% and bottom 25% in instantaneous velocities at each time, respectively. A coefficient significantly < 1 corresponds to high correlation, ≈ 1 corresponds to no correlation, and > 1 corresponds to anti-correlation. Control data is computed by random assignment of velocities to cells at each time (95% confidence interval of 100 simulations). Scale bars: 100 μm.

The longitudinal tracking data permitted various single-cell analyses and grouping in terms of behavioral phenotypes. We classified cells depending on their average motility (i.e. the ratio of travel distance to the tracking duration, see Supplementary Fig. 11). While the low-motility group (< 25%) cells were found to remain in the core of the spheroid, the trajectories of high-motility group (> 75%) cells feature outward migratory behavior and invasion into the surrounding gel matrix, particularly along paths near the glass plate (Fig. 6e). These two functionally distinct groups appeared to have originated from statistically distinctive regions in the tumor at early times (Supplementary Fig. 11). From the trajectory of each cell in every 6-hour window, the instantaneous velocity was calculated, and velocity maps at each time were obtained (Fig. 6f). The time-lapse video of the velocity maps shows how individual cells move during spheroid growth and invasion (Supplementary Video 6). Interestingly, the high-motility cells had consistently higher speeds than low-motility cells throughout the entire duration, and underwent moderate acceleration at 40–60 hours at the onset of invasion (Fig. 6g). In terms of instantaneous velocity, fast and slow cells (top and bottom quartile) were analyzed at each time point. A Geary’s coefficient analysis showed significantly higher spatial correlation of velocities among the fast cells compared to the slow cells and a randomized control, suggesting the presence of non-cell-autonomous behaviors^42^ (Fig. 6h). In particular, significant spatial correlation at earlier time points (20–40 hours) is attributed to the streams of cells moving in small packs within the spheroid (Supplementary Fig. 11). These results demonstrate the novel capability of gathering large-scale longitudinal single-cell information *in situ* using laser particles.

## Discussion

We have demonstrated that thickly-coated semiconductor microdisks are well suited for cell tagging and tracking applications due to their low pump energy, excellent stability, and biocompatibility. With respect to the spontaneous emission typical of fluorophores, the stimulated emission from single-mode microlasers has several distinct characteristics, such as coherent-state statistics, sharp threshold, directionality, and picosecond-scale decay times, that are potentially useful for imaging and sensing^38,39^. Most remarkably, the spectral width of laser emission is 100 times narrower than typical fluorophores. We harnessed this property for wavelength-encoded tagging and tracking of thousands of densely populated cells in a 3D scattering tissue.

Improvements of multiplexing capability are possible. First, the total wavelength span can be expanded to the NIR-I and visible range using appropriate III-V and II-VI semiconductor materials. Secondly, when multiple laser particles with different wavelengths are combined, the number of unique identifiers scales as 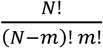 where *N* is the number of colors for singlet LPs (*m* = 1) and *m* is the number of particles in the multiplet. With *N* = 1000, doublets (*m* = 2) can have a half million identifiers, and the number increases to 166 million for triplets (*m* = 3). Preliminary results showed promising feasibility of this massively scalable approach (Supplementary Fig. 12).

Recent work has highlighted the importance of cellular heterogeneity at the single-cell level^40^. While current understanding primarily stems from sequencing of single dissociated cells, emerging spatial transcriptomic techniques promise understanding of cellular identity in the tissue context^41^. Laser-particle-enabled cell tagging could allow sorting and extraction of cells of interest for post-hoc analyses such as flow cytometry or single-cell sequencing^42^. Furthermore, it can enable highly multiplexed cell tracking of individual cells, offering rich complementary information on cellular behavior, including migration, motility, cell-cell interactions, and spatial clustering. The combination of imaging, single-cell tracking, transcriptomic and proteomic assays will enable unprecedentedly comprehensive evaluation of cell identity and function.

## Methods

### Fabrication

Microdisk resonators were fabricated starting from wafers composed of an InP substrate and two epitaxial layers: a 200-nm active layer of lattice-matched, undoped InGaAsP or InAlGaAs and a 100-nm capping layer of (undoped) InP. The capping layer was introduced to protect the active layer during the reactive ion etching (RIE) process.

For optical lithography samples, 2 μm of SU-8 photoresist (MicroChem) were spun coated on cleaned wafers and then soft baked (1 min at 65 °C, 2 min at 90 °C, 1 min at 65 °C). The mask used for photolithography (chrome on quartz, Benchmark Technologies) had a hexagonal pattern of holes (2.5 μm in diameter, spaced by 6 μm) with a density of 3.2 million microdisks per cm^2^. After the photoresist deposition, the wafer was diced into squares of 5 to 10 mm to produce about a million microdisks per batch. Photolithography consisted of exposure of the i- and h-line of a mercury arc lamp at a total dose of 40-60 mJ/cm^2^ (Karl Suss MJB4 mask aligner), a post-exposure bake at 90 °C, development in SU-8 Developer (MicroChem), and rinsing in isopropyl alcohol (IPA). A hard baking step at 200 °C was performed to harden the SU-8 pattern and smoothen its surface, followed by descum in O_2_ plasma (Anatech Barrel SCE 160). For dry etching, inductively-coupled-plasma reactive ion etching (ICP-RIE) (Unaxis Shuttleline) was conducted using a hydrogen bromide chemistry, which resulted in an etching depth of ~700 nm. After RIE, the photoresist was removed by mechanical scrubbing with cleanroom foam swabs, and the sample was rinsed in acetone, IPA and de-ionized (DI) water. A second O_2_ plasma treatment and an immersion in polymer stripper PRS-3000 (J.T. Baker) were performed to further remove residues.

For e-beam lithography samples, the semiconductor wafer was first coated with 330 nm of SiO_2_ by plasma-enhanced chemical vapor deposition (Orion III, Trion) to serve as a hard mask for the semiconductor etching process. Circular patterns of the desired dimensions were defined by 100 keV electron-beam lithography (JBX6300-FS, JEOL) on a negative-tone resist (ma-N 1410, MicroChem), and transferred to the silica hard mask by ICP-RIE (Plasmalab 100, Oxford Instruments) using fluoroform chemistry. A second etching step based on silicon-tetrachloride was used to transfer the pattern to the semiconductor. The remaining SiO_2_ hard mask was removed with a final fluoroform-based RIE step.

### Transfer and silica coating

For characterization in air, the post-RIE samples were partially wet-etched in a 3:1 solution of HCl in DI water for 5 to 10 s, leaving the microdisks suspended on a small pillar. For completely detaching the microdisks, the substrates were wet-etched face down in the HCl:H_2_O solution inside a 5 ml centrifuge tube for 30 s. The suspension of microdisks was then transferred to a 1 μm pore centrifuge filter, and filtered thoroughly by at least 3 repeated cycles of centrifugation and resuspension (via ultrasonication) using ultrapure water or ethanol (EtOH). The sample collection efficiency transferring from substrate to solution was typically around 50%.

Silica coating of the microdisks was performed by multiple cycles of a modified Stöber process. In a typical reaction cycle, microdisks (~10^6^ disks/ml) were suspended in 670 μl of an EtOH:H_2_O solution (80 v/v% EtOH). Next, 60 μL of 40 mM tetraethyl orthosilicate (TEOS) in ethanol, and 45 μL of ammonium hydroxide solution (28 v/v% NH4OH) were added, and the microdisk solution was shaken vigorously at 1400 rpm for 6 h at room temperature. To harden the silica shell to improve chemical stability, the temperature was increased to 80°C and the solution was kept mixing for at least 12 h^43^. Following each coating reaction, the microdisks were thoroughly centrifuge filtered and resuspended in water or ethanol. Multiple cycles of coating results in occasional (<10%) formation of laser particle multiplet aggregates. To fabricate a fluorescein-doped silica shell, fluorescein isothiocyanate (FITC) was reacted with 3-aminopropyl-trimethoxysilane (APTMS) and the resulting conjugate, FITC-APTMS, was used instead of TEOS in the coating reaction.

Surface functionalization was conducted by reacting coated microdisks with silane reagents. Coated microdisks in 600 μl of water (~10^5^ μdisks/ml) were mixed with 60 μl of 20 mM silane-polyethylene glycol-FITC (5 kDa) or silane-polyethylene glycol-biotin (5 kDa) in water, 6 μl of 20 mM TEOS in ethanol, and 4 μl of ammonium hydroxide solution (28 v/v% NH_4_OH). The solution was mixed at 1400 rpm at 70°C for 3 h, and then kept mixing at room temperature for at least 12 h, before centrifuge filtration and resuspension.

Samples embedded in 3D hydrogel matrix was prepared by mixing equal volumes of an aqueous solution of microdisk laser particles (coated or uncoated) with Matrigel (Corning) and incubating for 2 h at 37 °C to allow matrix cross-linking. For cell experiments, laser particles were resuspended in sterile filtered DI water before adding to cells.

### Modeling of microdisk resonance

Theoretical modeling of uncoated microdisk resonances for different diameters and external refractive indexes were performed as described in Supplementary Note 1 with a custom MATLAB script. Modeling of microdisk resonance sensitivity for different silica coating thicknesses was conducted via a series of fully-3D finite-difference time-domain simulations (Lumerical FDTD solutions). The semiconductor microdisk was modelled as a cylinder of refractive index of 3.445, while the refractive index of the silica shell was taken from tabulated values (Handbook of Optical Constants of Solids I–III by E. Palik). The simulation region was set with perfectly-matched layer (PML) boundary conditions on all directions. The distance to the PML boundaries as well as the meshing size were chosen after a series of convergence tests. The optical modes supported by the resonator were excited by a number of randomly-placed in-plane dipole emitters, and the resonance wavelength at the bandwidth of interest was recorded as the background refractive index of the simulation was varied.

### Electron microscopy

10 μl of laser particle suspensions in water were added and air-dried on silicon wafer chips for scanning electron microscopy (SEM), and formvar-carbon coated nickel mesh grids (Electron Microscopy Sciences) for transmission electron microscopy (TEM). TEM images were obtained using a JEOL JEM 1011 transmission electron microscope at 80 kV. SEM characterization was performed on a Hitachi S-4800 and a Zeiss Ultra Plus Field-Emission microscopes at 2 keV. For cross-sectional viewing, coated microdisks were first milled with a focused Ga^+^ beam using a dual-beam SEM/FIB tool (Helios Nanolab, FEI Company). Energy-dispersive X-ray spectroscopy and mapping was performed on a Zeiss Supra55VP Field Emission microscope at 8 keV.

### Optical characterization

For optical characterizations and imaging of microdisks, a commercial laser-scanning confocal microscope (Olympus FV3000) was modified. A pump laser (Spectra Physics VGEN-ISP-POD, 1060-1070 nm, pulse duration 3 ns, repetition rate 2 MHz) was coupled to a side port of the laser-scanning unit of the microscope. The emission from microdisks was collected from the same port and relayed by a dichroic mirror to a NIR spectrometer using an InGaAs linescan camera (Sensor Unlimited 2048L), operated with an integration time of 1 ms for high-resolution characterization; for LASE imaging, the acquisition of spectrometer data was performed at 100 μs integration and synchronized to the laser-scanning unit. A 600 lines/mm grating was used for high-resolution characterization (0.2-nm resolution, 150-nm span). For LASE imaging, a 200 lines/mm grating (0.6-nm resolution over 1150-1600 nm) was used. An NIR-optimized, 20X, 0.45-NA objective (Olympus IMS LCPLN20XIR) was used for LASE microscopy. Confocal fluorescence images were acquired with a 40X, 0.7-NA objective (Olympus LUCPLFLN40X). A stage-top incubator (Tokai Hit) kept the samples at 37 °C and 5% CO_2_ during imaging.

Stability of laser particle emission was performed in air for uncoated microdisks and in hydrogel for both coated and uncoated microdisks. For continuous pumping, individually pumped with *E_p_* = 40 pJ/pulse for 15 min, while collecting spectra every 15 s. For long-term stability, emission was collected via LASE imaging every hour for 30 h. The output intensity and wavelength were determined from the area under the spectral curve and the Gaussian-fit center respectively.

Sensitivity of laser particle’s emission to external refractive index was measured in different sucrose solutions prepared with concentrations of 0, 20, 38, 50, and 64% w/w. Their refractive index was measured using a portable refractometer (PAL-RI, ATAGO). Uncoated and coated (150 nm) laser particles were deposited on different wells of a glass-bottom 96-well plate. For each concentration, 300 μl of sucrose solution was pipetted into each well and a LASE image was acquired. Between successive measurements, the wells were washed twice with ultra-pure water. Note that under this experimental arrangement, only the refractive index of the top half-space is varied, while the bottom half remains constant throughout (glass).

### Cell culture and biocompatibility experiments

HeLa human cervical cancer cells (ATCC), 4T1 mouse breast tumor cells (ATCC), MDCK-II canine kidney epithelial cells (ECACC) were cultured and maintained in serum-supplemented cell media following manufacturer guidelines. Membrane-GFP expressing HEK-293 human embryonic kidney cells were a gift from Prof. Adam E. Cohen, and were cultured in Dulbecco’s modified Eagle medium (DMEM) supplemented with 10% (v/v) fetal bovine serum (FBS) and 1% (v/v) penicillin-streptomycin. Cells were stained after fixation with AlexaFluor 594-Phalloidin for actin (Thermo Fisher), and DAPI for nucleus (ProLong Gold AntiFade Mountant with DAPI, Thermo Fisher) following manufacturer guidelines.

A typical protocol to load laser particles into cells is as follows. Cells were plated in their respective media at a known density in a glass-bottom, 96-wells plate. Laser particles were resuspended in sterile filtered DI water, and counted using a standard hemocytometer. The laser particle solution was then added to cells, at initial particle-to-cell ratios from 1:1 to 4:1. Immediately afterwards, a requisite amount of 10x PBS was added to maintain isotonicity. The dilution of cell media with addition of laser particle solution was <10%. The cell media was exchanged to fresh media within 2 h. Cells were then incubated at 37 °C and 5% CO_2_ for 24-48 h until laser particle uptake was complete.

Cellular viability was assessed via a calcein-AM/ethidium homodimer-1 fluorescent assay (LIVE/DEAD Viability/Cytotoxicity Kit, Thermo fisher). MDCK-II cells were switched to serum-free MEM-alpha for the duration of the experiment (up to 72 h) to minimize cell proliferation. Uncoated or coated microdisks in water were added to MDCK-II cells such that the particle-to-cell ratio was approximately 1:1. Fluorescent staining was conducted after 24, 48 and 72 h later. Fluorescence and bright-field microscopy was conducted to quantify particle uptake and cell viability (Keyence BZ-X700 microscope).

Cellular proliferation was assessed via a cell counting kit-8 (CCK, Millipore Sigma). After MDCK-II cells were plated, low (4:1 particle-to-cell) and high (32:1) concentrations of microdisks were added as previously described. CCK-8 staining was conducted at 48 and 120 h, when cells were approximately 80% confluent. Absorbance measurements were taken using a spectrophotometer (Epoch 2, BioTek Instruments).

### Time-lapse bright-field imaging

To observe the internalization and interaction of cells with laser particles, we used an automated, time-lapse, bright-field microscope (Keyence BZ-X700). MDCK-II cells were plated with laser particles (2:1 particle:cell ratio) in a 96 well glass-bottom plate as described earlier. Images from different wells were taken every 5 min over a 60 h period in a stage-top incubator (Tokai Hit) at 37 °C and 5% CO_2_. After correcting for drift artifacts, the processed videos were analyzed frame-by-frame to record instances of particle uptake, cell division, and suspected exocytosis events. Over 400 individual cell cycles were annotated, from which we calculated the cell cycle time as a function of particle number, and the statistics of particle partitioning due to cell division. Empirical probabilities for laser particle partitioning were fit with an asymmetric binomial distribution: *P*(*r*) = *C*(*n,r*)*p^r^*(1 – *p*)^*n*–*r*^, where *n* is the number of particles in the parent cell, *r* is the number of particles in the daughter cell, *C*(*n,r*) = 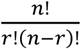, and *n* is a fitting parameter that quantifies the asymmetry of the distribution. *n*=0.5 for a symmetric binomial distribution.

### *Ex vivo* lung imaging

The MGH Institutional Animal Care and Use Committee approved our animal protocol (2017N000021) in accordance with NIH guidelines. Membrane-GFP expressing mTmG mice were purchased from Jackson Laboratories. 4T1 cells were loaded with laser particles *in vitro* as previously described, and stained with CellTracker Red (Thermo Fisher) following manufacturer guidelines. Approximately 2x10^5^ 4T1 cells containing ~1x10^5^ laser particles in 200 μL of DPBS solution was injected intravenously into the tail vein of a membrane-GFP expressing adult mouse under anesthesia. After 30 min, the animal was euthanized, and the lung was explanted. The entire lung was immersed in DPBS and placed on a glass-bottom dish. A 60x, 1.2-NA objective (Olympus UPLSAPO 60XW) was used for confocal fluorescence and LASE imaging up to 150 μm in depth. For large-area imaging, 64 adjacent areas, each spanning 212 by 212 by 3 μm^3^ were imaged and stitched together in post-processing using Fiji and MATLAB.

### Spheroid invasion assay

Approximately 1.5x10^5^ laser particles were added to 5x10^3^ 4T1 cells as previously described and incubated for 48 h until confluent in a 96-well cell culture plate. Cells were transferred to an ultra-low attachment (ULA)-coated round-bottom microplate (CellCarrier) to form spheroids. Each well initially contained approximately 2x10^4^ cells and 7x10^4^ laser particles. After 3 days of incubation, the spheroids were transferred to a glass-bottom microplate in 100 μl of serum-supplemented cell media mixed with 100 μl of Matrigel (Corning). After 2 h, an additional 100 μl of cell media was added. The spheroids were used for LASE imaging after 6 h. The spheroid was imaged every hour for a total of 129 measurements (*t* = 0, 1, …,128 h). The imaged region was divided in 4 adjacent areas acquired sequentially; each area was 480 by 480 by 280 μm^3^ and was imaged at 320 by 320 by 70 pixels, 100 μs/pixel; total imaging time for each scan was ~48 mins (~12 mins per area). Laser pumping was set at *E_p_* = 80 pJ/pulse.

### LASE imaging data analysis

A custom Python software was used to analyze the spectra acquired during LASE imaging in real time, recording the data only when they contained peaks greater than a threshold level. Post processing of the data was performed with a MATLAB custom code. The recorded spectra were further reduced to wavelength peaks by Gaussian fitting, producing a set of wavelength (*λ*) and position 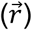 data of each recorded “lasing” event (or pixel). Since many lasing events can be recorded from a single laser particle, a clustering algorithm was applied to group the data from the same laser particle. A three-step hierarchical clustering algorithm was used: (i) clustering by position to isolate spatially separate clusters of lasing pixels; (ii) clustering by wavelength to distinguish any adjacent particles with different wavelengths; (iii) clustering by position again to separate clusters in the cases where two particles with similar wavelengths were bridged by a third particle with a different emission. Once clusters are identified, their centroid position and mean wavelength were computed to represent corresponding laser particles.

For particle-in-cell tracking in tumor spheroids, a metric (*D_ij_*) was introduced, which represents a “distance” between two particles identified in two separate measurements in 4-dimensional space of wavelength (*λ*) and position 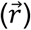; the distance *D_ij_* between the *i*-th particle in an image and the *j*-th particle in a preceding image is defined as: *D_ij_* = 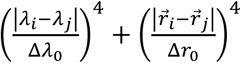, where Δ*λ*_0_ and Δ*r*_0_ are adjustable parameters and set to 1 nm and 20 μm for optimal results. For each time point *t_m_*, the distance *D_ij_* of all imaged particles (*i* = 1 to *N_m_* from those (*j* = 1 to *N_m-1_*) detected in the previous time (*t_m-1_*) were calculated. Pairs of particles with a distance smaller than a threshold *D_th_* = 2 were matched, starting from the one with the smallest distance. For the remaining particles, the matching procedure was repeated with the particles detected at earlier times (*t_m-2_*, *t_m-3_*, and *t_m-4_*). Laser particles within the same cell were identified by computing for each couple *i, j* of tracked particles their average (geometric) distance *S_ij_* = 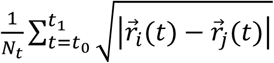 at each time point. Particles with *s̅_ij_* < 21 μm were considered belonging to the same cell for the entire duration of the tracking and were thus merged in a single trace.

To remove the contribution of intracellular movements of laser particles on the overall cellular trajectories, we applied a 6 hour moving average to the tracked pathways.

### Statistics

Data were presented as either box and whisker plots, or as mean and 95% confidence interval. Statistical hypothesis testing for the biocompatibility assays was done using either one-way or two-way ANOVA, and all F-statistics were reported. Statistical significance was set at *α* = 0.05. Post-hoc comparisons were conducted using Tukey’s test. Statistical analyses were performed using GraphPad Prism and MATLAB.

To quantify spatial autocorrelation in the spheroid tracking experiment, we computed Geary’s coefficient as follows: *C* = 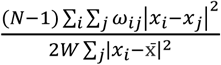, where *N* is the number of neighboring cells, and *ω_ij_* is a spatial weight matrix that defines the neighborhood region of interest, or the pack size of spatially correlated cells. Before normalization, *ω_ij_* contains zeros on the diagonals and ones for neighboring cells within a given distance. This distance is set to 60 μm in Fig. 5g, and is varied from 30 μm to 600 μm in Extended Fig. 7g-h. The matrix is then row-normalized such that each row sums to 1. *W* is the sum of all *ω_ij_* elements. *x_i_* and *x_j_* are either the (vector) velocities of cell *i* or *j* at a given time-point (Fig. 5g) or the overall motility (displacement / time-tracked) of cell *i* or *j* (Extended Data Fig. 7g). x̄ is either the mean velocity at a given time-point or the mean motility. To generate a control sample, the Geary coefficient was also computed for cells with corresponding cell velocities or motilities randomized. The randomization process was repeated 100 times, and the 95% confidence interval of the random control group was presented.

## Acknowledgments

This work was supported by the U.S. National Institutes of Health grants DP1-OD022296, P41-EB015903, R01-CA192878, and National Science Foundation grants ECCS-1505569. Electron microscopy was performed in the Microscopy Core of the Center for Systems Biology/Program in Membrane Biology, which is partially supported by an Inflammatory Bowel Disease Grant DK043351 and a Boston Area Diabetes and Endocrinology Research Center (BADERC) Award DK057521. This research used resources of the Center for Functional Nanomaterials, which is a U.S. DOE Office of Science User Facility, at Brookhaven National Laboratory under contract (DE-SC0012704) and of the Center for Nanoscale Systems, a member of NNCI, which is supported by the NSF under award no. 1541959.

The authors would like to thank Fernando Camino for helping with FIB measurements.

